# Structural insights into viral hijacking of p53 by E6 and E6AP

**DOI:** 10.1101/2023.11.01.565136

**Authors:** Colby R. Sandate, Deyasini Chakraborty, Lukas Kater, Georg Kempf, Nicolas H. Thomä

**Author notes:** Authors contributed equally (Colby R. Sandate, Deyasini Chakraborty).

## Abstract

The E3-ubiquitin ligase E6AP degrades p53 when complexed with the viral protein E6 from human papilloma virus (HPV), which contributes to the transformation of cells in HPV-related cancers. Previous crystal structures of the E6AP-E6-p53 ternary complex have implicated a peptide containing an LxxLL motif from E6AP as the interface between the three proteins. However, the contributions to the ternary complex from the remainder of the E6AP protein remain unknown. We reexamined this complex using cryo-EM and full-length proteins and find additional protein interaction interfaces involving a previously uncharacterized domain of E6AP. Additionally, we observe that the ternary complex forms both 1:1:1 and 2:2:2 stochiometric complexes comprised of E6AP, E6 and p53.

## Introduction

p53 is a multi-domain transcription factor known for its role as a tumor suppressor protein and was originally shown to arrest the cell cycle at G1 phase in response to DNA damage^1^. It is now known that p53 performs a large variety of additional functions both related and unrelated to tumor-suppression which are cell-type or cell-state dependent^2^.

The polypeptide chain of the p53 protein is subjected to numerous post translational modifications (PTMs). Depending on the type and location of the modified residue, these PTMs can regulate p53 levels, cellular localization, and function^3^. Interestingly, covalent modification of p53 by ubiquitin (Ub) has been reported to affect all three of these modes of regulation. Protein levels are regulated in large part by the ubiquitin proteasome system (UPS), involving various ubiquitin-related enzymes which polyubiquitinate proteins targeted for proteolytic degradation via the proteasome^4^. This is also the case for p53, which is polyubiquitinated by the E3 ligase MDM2 in addition to others^3,5^.

While p53 is normally regulated by the UPS, it has also been shown to be a common target of pathogenic viruses, which affect the cell cycle with the help of the UPS. Indeed, p53 was originally discovered as a 53 kDa host protein bound to the oncogenic simian virus 40 large T antigen^6,7^. Since that initial discovery, many more viral proteins have been found to directly interact with p53, including the E6 protein from the Human papilloma virus (HPV). HPV is a nonenveloped double-stranded DNA virus which has been identified as the etiological agent of various human carcinomas, as well as causing benign lesions forming warts in the genital tissues and in the upper respiratory tract^8,9^. While prophylactic HPV vaccines have been available since 2006, only a minority of at-risk individuals have received them^10^. New treatment options are needed, as prophylactic vaccines do not eliminate persistent HPV infection or inhibit progression^11^.

The oncogenicity of malignant variants of HPV is achieved via expression of viral proteins E6 and E7 in the infected cell, which degrade host proteins p53 and retinoblastoma protein, respectively^8,9^. E6 degrades p53 through direct interaction and subsequent recruitment of the E3 ligase E6-associated protein (E6AP, also known as UBE3A) to p53 in a ternary complex^12^. E6AP is a founding member of the HECT ligases, and unlike the RING E3 ligases, has intrinsic catalytic activity and forms a Ub-conjugated thioester intermediate prior to transfer of Ub to the target protein^4^. E6AP has been found to not only be important for normal neurological development, but to also require a precise dosage in the cell for normal function. Duplications and deletion in the maternally inherited allele as well as loss-of-function mutations of E6AP have been shown to produce Angelman Syndrome and Angelman Syndrome-like symptoms while gene duplication is also associated with Autism Spectrum Disorder in patients^13–15^.

E6AP consists of multiple structured domains. Near the N-terminus (aa 24-87) is a small zinc-binding AZUL domain, consisting of a helix-loop-helix motif which binds hRPN10 in the proteasome lid, with an unknown function^16,17^. At the C-terminus is a HECT domain (aa 518-875), divided into a N- and C-lobe. The C-lobe contains the catalytic residue C820, while the N-lobe contains the binding site for E2 enzyme UbcH7^18^. A previous study crystallized the ternary complex of E6AP-E6-p53, using a 12 residue peptide of E6AP containing a LxxLL motif (aa 403-414) and found that it forms the core of the ternary interaction^19^. This peptide inserts into a pocket of E6, which then structures a p53-interacting cleft. While the HECT domain^18^ and the LxxLL motif^19^ have each been separately crystallized, is unknown how these separate structural elements are oriented with respect to one another in the E6AP’s full strucrure. Additionally, it remains to be seen if other parts of the E6AP protein besides the 403-414 LxxLL motif are involved in the interaction comprising the ternary complex.

## Results

Building on previous work analyzing the ternary complex formed between the HPV viral E6 protein, E6AP and p53 via crystallography, we sought to revisit the structural characterization using cryo-EM with full-length proteins in the cellular context of HPV infection. Using recombinantly purified proteins, we were able to show complex formation using mass photometry (MP) on a equimolar mixture of E6AP, E6 and full-length p53 (**Supplementary Fig. 1A**). The MP results revealed the prescence of two populations at 507 +/− 43 and 636 +/− 68 kDa in mass. These are considerably larger than the calculated mass for a 1:1:1 stochiometric complex (176 kDa) or even a p53 tetramer bound to a single copy of E6AP and E6 (222 kDa). In addition to the two larger populations, we also observed a population at 105 +/− 20 kDa and another at 167 +/− 35 kDa, which are similar to calculated masses of monomeric and dimeric E6AP (100 and 200 kDa, respectively). These populations were also in agreement of MP measurements of E6AP alone (**Supplementary Fig. 1B**), which showed a major population at 88 +/− 12 kDa (monomer) and a minor population at 172 +/− 27 kDa (dimer). We also analyzed this protein mixture at higher concentration by SEC-MALS, an orthogonal mass determination technique that allowed for higher sample concentration compared to MP, in case the lower sample concentration used in MP (40 and 20 nM) did not allow for complete complex formation. The SEC-MALS measurement agreed with the MP results and also showed the formation of larger complexes, where three peaks were observed with a calculated mass of 671 +/− 175 kDa, 462 +/− 167 kDa, and 221 +/− 74 kDa (**Supplementary Fig. 1C, Supplemenatry T2**). Overlaying a separate run with isolated p53 shows a single peak with a calculated mass of 219 +/− 59 kDa that elutes at a different volume compared to the peaks observed in the ternary complex run. To test whether the DNA binding domain (DBD) of p53 was sufficient to drive complex formation, we also analyzed a equimolar mixture of E6AP, E6 and a truncation of p53 containing only the DBD (p53^DBD^) by MP (**Supplementary Fig. 1D**). We again see two populations likely corresponding to monomeric and dimeric E6AP (129 +/− 22 kDa and 193 +/− 25 kDa, respectively). We also observe a new population at 263 +/− 36kDa which is too large for a 1:1:1 stoichiometric ternary complex but is smaller than those observed in complexes containing full-length p53.

We next subjected the ternary complex to gradient fixation (graFIX)^20^ using glutaraldehyde (GA) or bissulfosuccinimidyl suberate (BS3) as a crosslinking reagent, prior to vitrification for analysis by single particle cryo-EM. The BS3 sample yielded a structure of the ternary complex at ∼3.7 Å global resolution (**Fig. 1A**) as measured by a gold standard Fourier Shell Correlation (FSC) of 0.143. Using rigid-body fitting, we were able to manually position the crystal structure (PDB:4XR8)^19^ of a single p53^DBD^ bound to E6 within the density. We were able to fit the majority of an AlphaFold2 predicted structure^21,22^ of full-length E6AP into the remaining unassigned density, with the exception of aa residues 1-120, which accounted for the N-terminus and AZUL domain.

**Figure 1.**
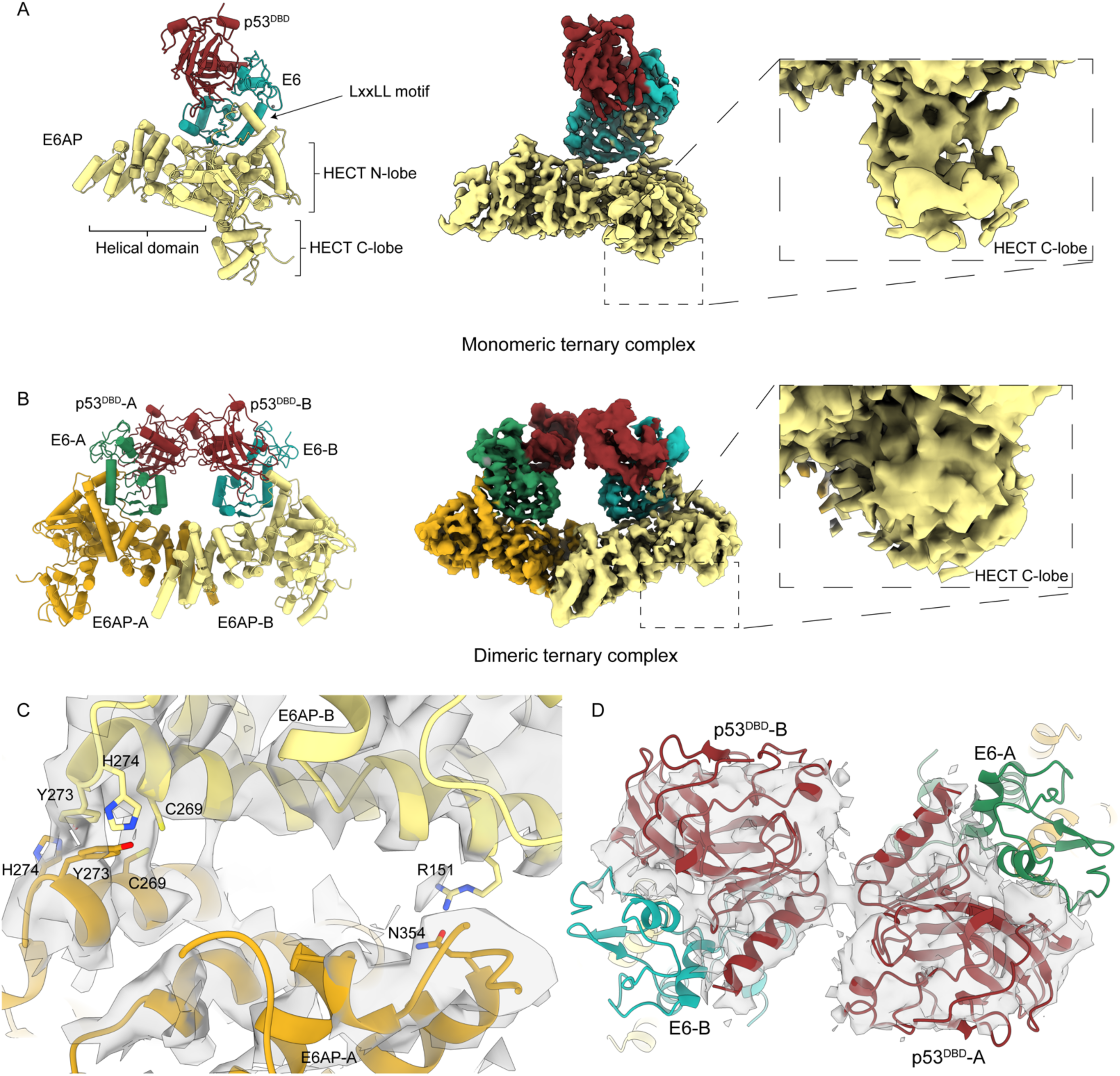
**A**) Structure of the monomeric ternary complex of E6AP-E6-p53. E6AP is depicted in yellow, E6 in green, and p53 in red. Left) Atomic model of the ternary complex in cartoon representation. Middle) Cryo-EM density color-zoned according to atomic model at high contour value. Inset) Cryo-EM density color-zoned according to atomic model at low contour value focused on the C-lobe of the HECT domain. **B**) Structure of the dimeric ternary complex of E6AP-E6-p53. The two protomers of E6AP are depicted in yellow and dark yellow, the two E6 protomers in light and dark green, and both protomers of p53 are shown in red. Left) Atomic model of the dimeric ternary complex in cartoon representation. Middle) Cryo-EM density color-zoned according to atomic model at high contour value. Inset) Cryo-EM density color-zoned according to atomic model at low contour value focused on the C-lobe of the HECT domain for one E6AP protomer. **C**) Zoom of E6AP dimer interface. Sidechains of residues likely contributing to the interface are shown in stick representation. Cryo-EM density is shown as semi-transparent. **D**) Zoom of p53^DBD^ dimer interface in dimeric ternary complex. Cryo-EM density is zoned around the two p53 protomers and shown as semi-transparent.

The cryo-EM dataset for the GA crosslinked sample resulted in a C_2_-symmetric map at ∼4 Å global resolution (**Fig. 1B**). In contrast to the BS3 sample, we observed density for “a dimer of ternary complexes” where two copies each of E6AP, E6 and p53 fit within the cryo-EM map. The main dimer interface composed of the E6AP N-terminal alpha-helices (**Fig. 1C**) and a minor interface formed by the two copies of p53-DBD (**Fig. 1D**), which together resemble the shape of a closed ring (**Fig. 1B**). Two copies of the monomeric ternary complex were used to fit within the density, without any notable structural changes outside of the dimerization of the ternary complex in comparison with the monomeric model. Interestingly, the p53^DBD^ dimer in the dimeric model shows a ∼35-degree rotation between the two protomers in comparison to crystal structures of a DNA-bound p53^DBD^ dimer^23,24^, or a single dimer within a DNA-bound tetramer (**Supplementary Fig. 2**)^25^. We then processed an additional BS3 dataset with increased ice thickness, to search for larger particles that would correspond to the dimeric complex seen in the GA dataset. Upon further analysis of the reference-free 2D classification results, we found a subset of particles that clearly resemble the dimeric ternary complex (**Supplementary Fig. 3**). This result, combined with the MP analysis of the in-solution complex showing E6AP monomers, dimers and large ternary complexes; suggests that the ternary complex is in an equilibrium between the monomeric and dimeric complex (and possibly larger, more transient species). Our workflow may have enrichened for the monomeric complex with BS3 and the dimeric complex with GA, based on the shorter and longer crosslinking diameter of the reagent, respectively.

In our structure, the N- and C-lobes of the HECT domain are clearly ordered and assume the same L-conformation previously observed in earlier crystal structures of the HECT domain^18^. The crystal structure of E6AP-E6-p53 contained a peptide from E6AP, corresponding to the LxxLL motif^19^, bound within a pocket formed between the N- and C-terminal zinc binding domains of E6 (**Fig. 1A, 2B**). Our structure also features the loop-helix-loop corresponding to this motif, however it deviated slightly from the published crystal structure and undergoes a large rearrangement compared to the AlphaFold2 monomer prediction of E6AP alone (**Fig. 2A**). Multimer prediction with AlphaFold2, which included E6 and p53, was able to predict the extended LxxLL conformation, but failed to predict the E6-p53 interaction and interactions between p53 and E6AP. Our structure also shows an additional interaction between a basic loop of E6AP near the LxxLL helix (414-418) and the 224-229 loop and R110 of p53. Similar to the LxxLL motif, this region of E6AP also undergoes a rearrangement when compared to the predicted apo structure, likely induced by binding to E6.

**Figure 2.**
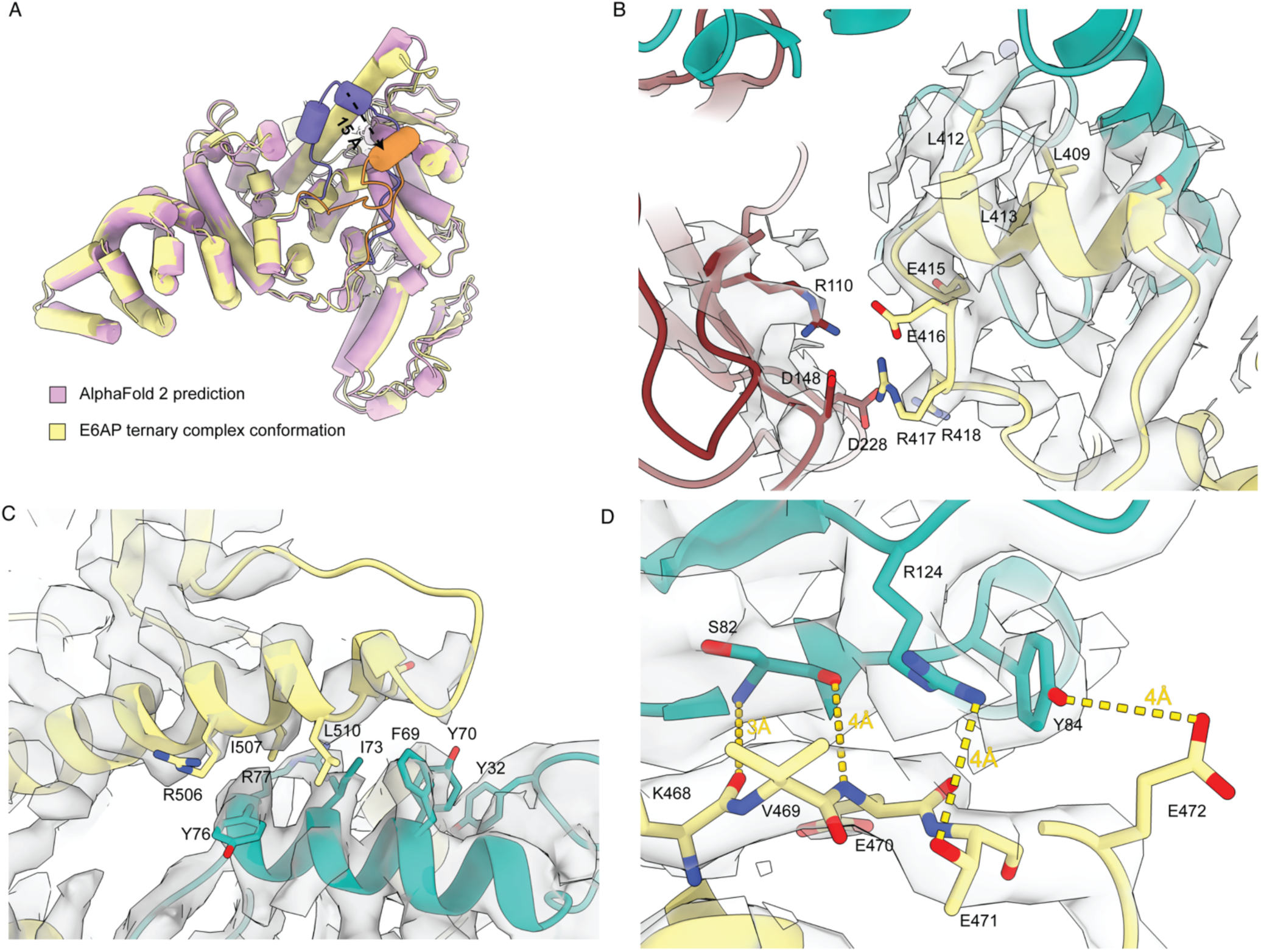
EPI domain of E6AP forms additional interfaces with p53 and E6. **A**) Superposition of AlphaFold2 monomer predicted structure of E6AP (pink) and the E6AP conformation solved while in the ternary complex (yellow). The loop-helix-loop pertaining to the LxxLL motif of E6AP is colored purple for the AlphaFold2 monomer prediction model and orange for the ternary complex conformation. **B**) Zoom of the LxxLL helix of E6AP embedded within the hydrophobic pocket of E6 and adjacent to p53. Residues are shown in stick atomic depiction and cryo-EM density is semi-transparent. **C**) Zoom of hydrophobic interactions between the 486-514 helix of E6AP and 64-77 helix of E6. **D**) Mainchain and sidechain interactions between the 468-476 loop of E6AP with S82, Y84 and R124 of E6. Possible H-bonds are denoted by dotted line with distances labelled in Å.

A large ordered domain consisting primarily of helices and comprising residues 130-516 is now resolved in our structure, which was previously undetermined by empirical methods but matches the AlphaFold2 predictions. Analysis of the domain by FoldSeek^26^ showed strong structural alignment only with other E6AP/UBE3a homologs and HECT-E3 ligases. This domain forms two additional interaction interfaces with the E6 protein outside of the LxxLL motif. The 486-514 helix of E6AP forms a hydrophobic interface with the 64-77 helix of E6 (**Fig. 2C**) on the opposite face of the hydrophobic cleft that binds the LxxLL motif of E6AP. Nearby, mainchain and sidechain interactions are present between the 468-476 loop of E6AP with S82, Y84 and R124 of E6 (**Fig. 2D**). Due to the functional nature of this domain in forming interactions with p53 and E6, we are designating it as the <u>E</u>6-<u>p</u>53 Interacting (EPI) domain.

## Discussion

Our structural characterization of full length E6AP in the ternary complex reveals three additional protein-protein interaction interfaces between the ligase and E6 and p53. Each of these interfaces belong to the previously uncharacterized EPI domain, which appears unique to E6AP homologs or highly similar HECT E3 ligases in other organisms. Additionally, our structure provides the first structural positioning of the LxxLL motif relative to the rest of the protein.

In both of our structures, p53 is distal from the catalytic HECT C-lobe, and it remains to be seen how this conformation results in Ub transfer from E6AP to p53. It has been proposed that the entire HECT domain may be flexible and allow for a rearrangement of the HECT C-lobe relative to the N-lobe that could bridge the distance between substrate and the catalytic cysteine^27–29^. We see that the C-lobe is flexible relative to the remainder of the complex in both our monomeric and dimeric ternary complex structures (**Fig. 1AB**, **Supplementary Fig. 3**-4). It is possible that our structures represent an ‘initial recognition’ or ‘collision complex’ between the ligase and substrate protein and that some other trigger is necessary for conformational change, such as co-binding the E2 enzyme, or the formation of a thioester Ub intermediate at the catalytic cysteine. Further work is necessary to test this hypothesis.

The question of E6AP’s oligomeric state has remained open since the publication of a trimeric structure of E6AP’s HECT domain^18,30^. Our data suggests that this oligomerization interface would result in a steric clash in the trimer due to the EPI domain, in the absence of a large-scale conformational change. However, our dimeric ternary complex suggests that E6AP can bind its p53 substrate as both a dimer and monomer in the presence of viral protein E6. Additionally, our full length E6AP protein purified from insect cells also shows evidence of a dimeric population in MP, even in the absence of other factors (**Supplemenatry Fig. 1B**). Upon addition of E6 and p53 in equimolar amounts, we can see the formation of higher order complexes, with measured masses that match expected values for a 3:3:3 and 4:4:4 stoichiometric complex (**Supplementary Fig. 1A**). This also agrees with the masses measured by SEC-MALS. Interestingly, when a truncation of p53 that only includes the DBD is included in the complex (**Supplementary Fig. 1D**), the largest mass we observe corresponds to the dimeric ternary complex previously seen in our cryo-EM data. This suggests that higher order oligomeric assemblies above a 2:2:2 stoichiometry are driven by other domains in p53 besides the DBD, such as the tetramerization domain. This is further supported by the relative orientation between the two protomers of the p53^DBD^ dimer present in our dimeric ternary complex structure (**Fig. 1D**). In comparison to crystallized tetramer structures of p53, where the DBDs are in a planar orientation to one another, there is a ∼35 degree rotation of one protomer outside of the planar orientation (**Supplementary Fig. 2**), which greatly reduces the contacts between these two protomers and likely inhibits the DBDs from assembling in the planar orientation necessary to build up the tetramer.

With the additional structured EPI domain resolved in our models, it is possible to reconsider the viral-induced ternary complex’s effects on p53 function. While it has been shown that p53 dimers can bind DNA and activate transcription^23,24^, the functional form of p53 is thought to be a tetramer of DBDs^25,31,32^. Superposition of our monomeric ternary complex upon one protomer of the p53 tetramer (**Fig. 3**) predicts several steric clashes between the adjacent protomer and E6 as well as the EPI domain of E6AP. We interpret this result to suggest that tetramer formation of the p53 domains may be disfavored while incorporated into the ternary complex, by preventing later contacts between protomers (such as between protomers A-B and C-D for example). However, dimers that are free of lateral contacts are unlikely to be affected (such as between protomers A-C and B-D) and may still be available to bind DNA.

**Figure 3.**
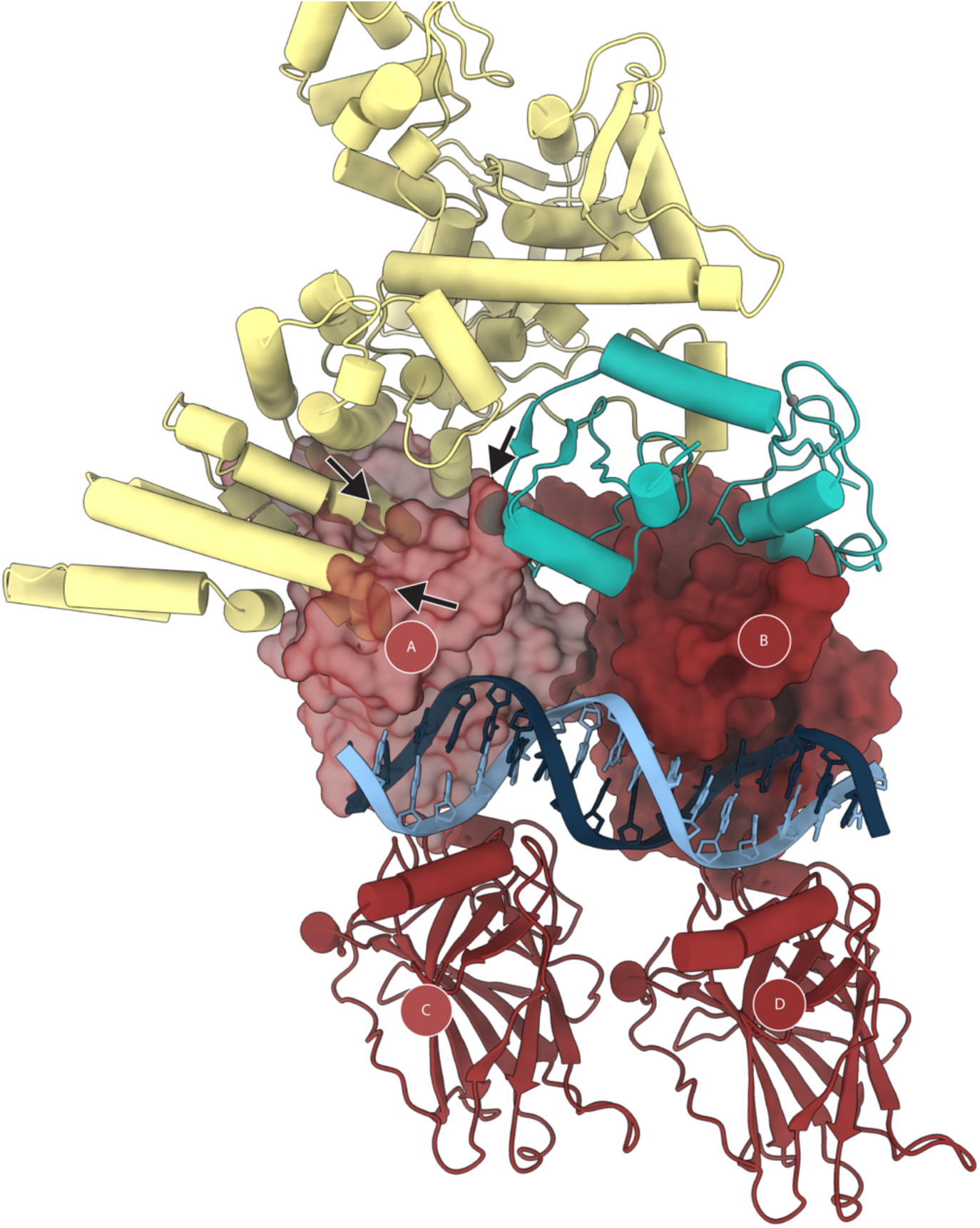
The tertiary complex produces steric clashes between p53 protomers in the DNA-binding tetramer. Superposition of E6AP-E6-bound p53 on a DNA-bound p53 crystal structure (PDB: 3KMD) shows the formation of steric clashes between adjacent p53 protomer A and E6 and E6AP (arrows). Protomers A and B are shown in surface representation, with Protomer A being semi-transparent.

To date, most functional studies have so far focused on the HECT domain of E6AP, and the region corresponding to the EPI domain has not been annotated for performing a particular function. Our structures suggest that the EPI domain is important fo E6AP dimerization, but it may also serve as a hub for interactions. Future work will be necessary to determine whether the EPI domain is involved in other protein and substrate interactions of E6AP, of which many others have been described^33–37^.

**Supplementary Figure 1.**
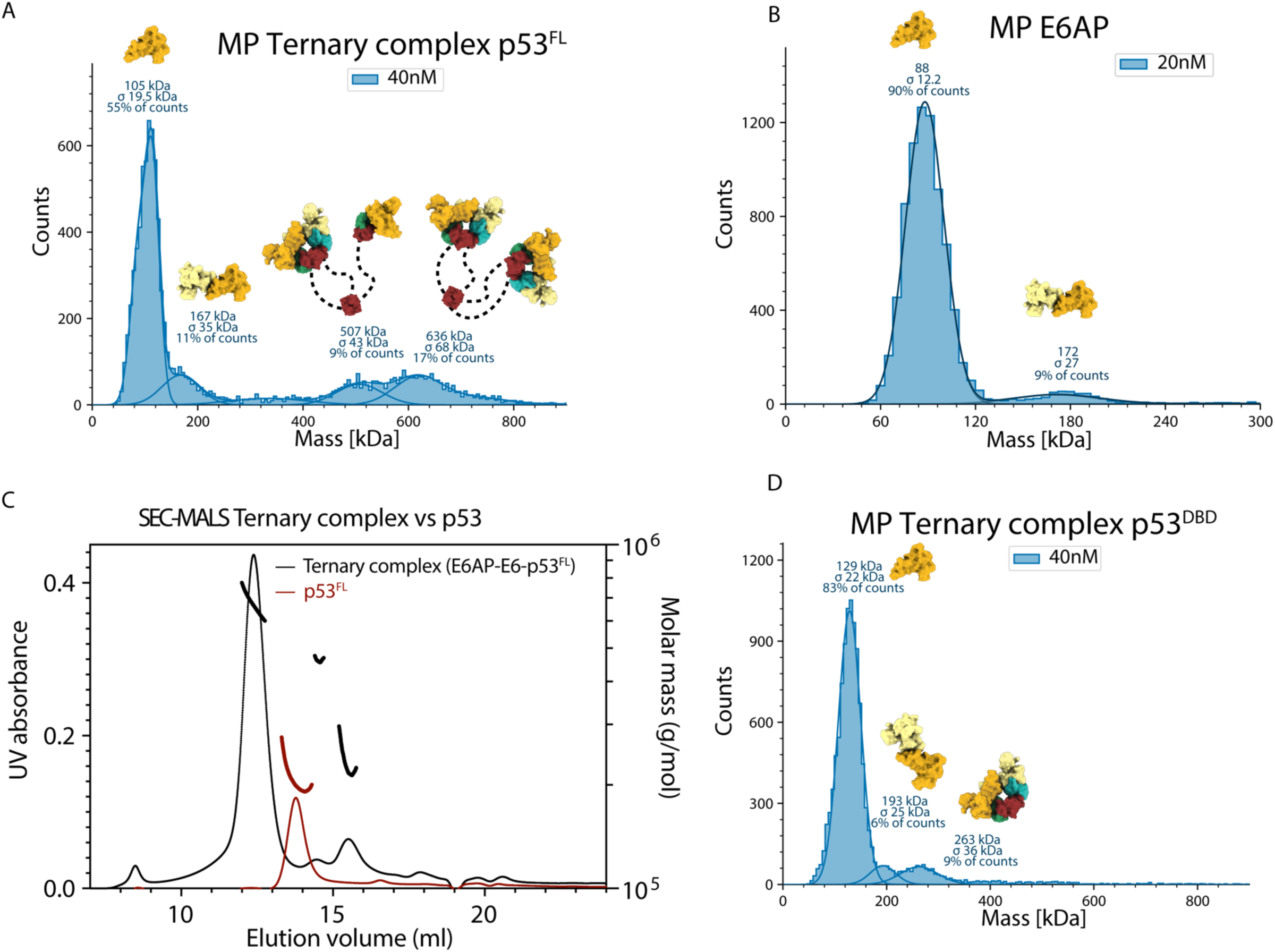
Biophysical characterization of fully formed ternary complexes of p53-E6-E6AP. **A**) MP mass distribution of 40 nM total concentration of full-length p53, E6 and E6AP proteins. Major bands were observed that corresponded to the mass of monomeric E6AP, dimeric E6AP, a 3:3:3 stoichiometric ratio of E6AP:E6:p53, and a 4:4:4 stoiciometric ratio of E6AP:E6:p53, respectively. **B**) MP mass distribution of E6AP, showing major bands corresponding to monomers and dimers. **C**) Size characterization of fully formed ternary complexes of E6AP-E6-p53 by SEC-MALS overlayed with p53^FL^ in isolation. **D**) MP mass distribution of 40 nM total concentration of p53^DBD^, E6 and E6AP. Major bands corresponded to the mass of monomeric E6AP, dimeric E6AP and a 2:2:2 stoichiometric ratio of E6AP:E6:p53^DBD^.

**Supplementary Figure 2.**
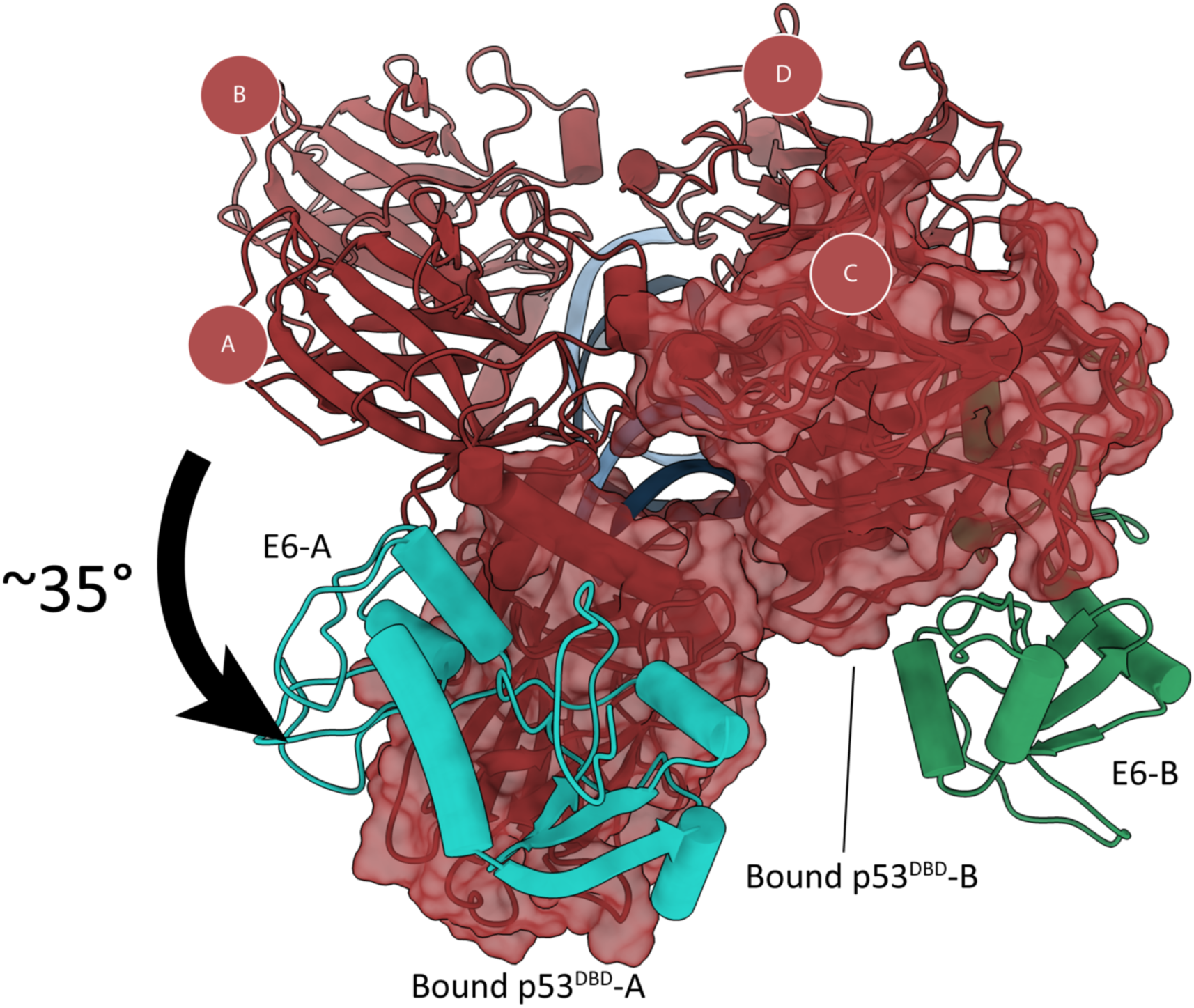
The p53 dimer bound in the dimeric ternary complex is shifted from it’s planar orientation within the DNA-binding tetramer. Superposition of p53 dimer (shown in semi-transparent surface representation) and bound E6 from dimeric ternary structure on Protomer C of the p53 tetramer crystal structure (PDB: 3KMD). Arrow depicts rotation of second bound protomer relative to Protomer A in the crystal structure.

**Supplementary Figure 3.**
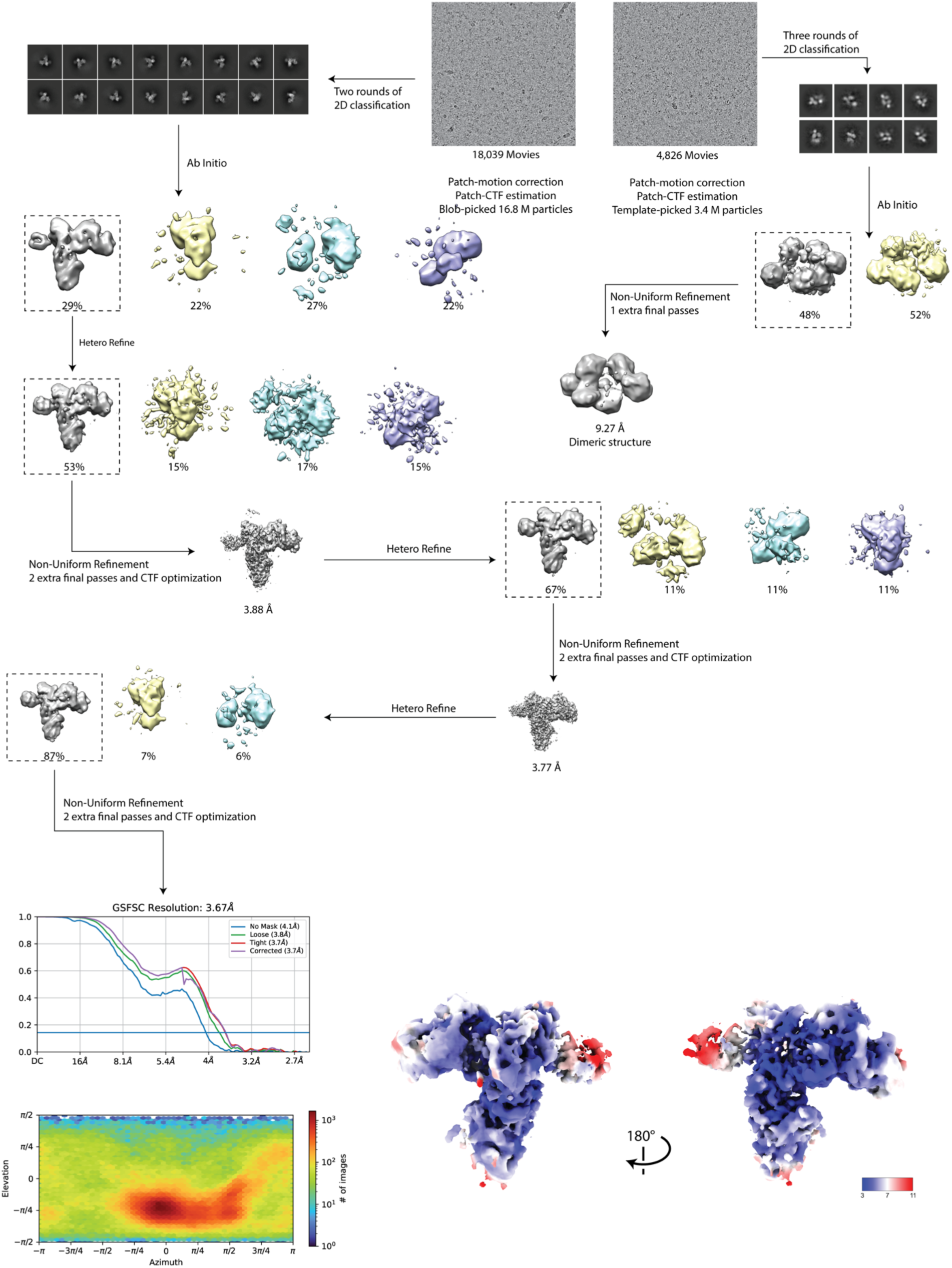
Cryo-EM data processing of monomeric E6AP-E6-p53 ternary complex. Image processing pipeline for monomeric ternary complex density maps and possibly dimeric complex maps derived from BS3 crosslinked datasets. All data were processed in cryoSPARC v.3 or v.4. Representative motion-corrected micrographs are shown at the top. Datasets were imported into cryoSPARC prior to motion correction and patch-based CTF estimation. Subsequent classification, ab inito and refinement steps are shown. The final model is colored by local resolution and presented in two orienatation, rotated 180 degrees. Viewing angle distribution and GSFSC curves are shown after FSC auto-tightening for the final monomeric ternary complex density map.

**Supplementary Figure 4.**
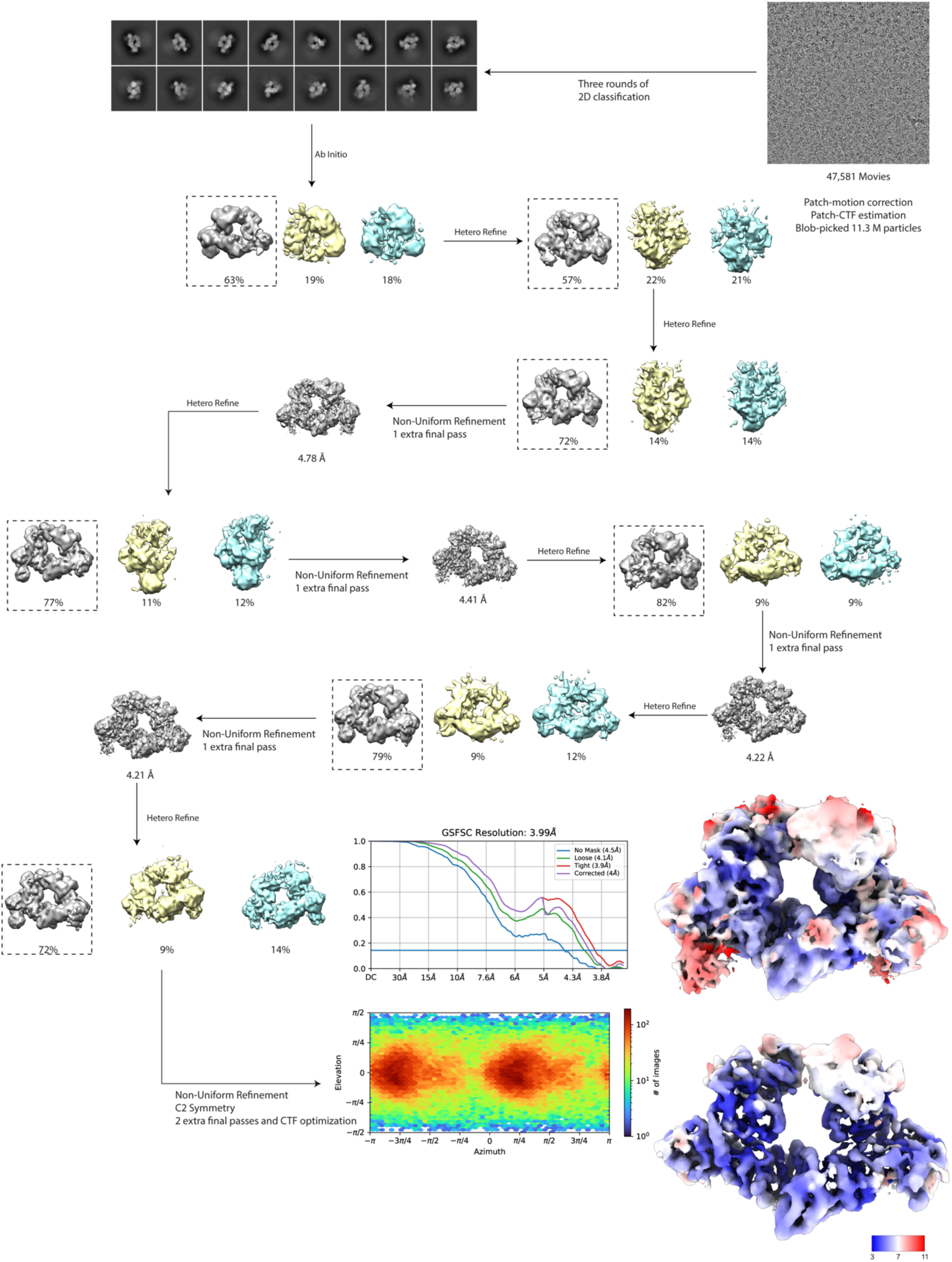
Cryo-EM data processing of dimeric E6AP-E6-p53 ternary complex. Image processing pipeline for dimeric ternary complex density maps derived from GA crosslinked datasets. All data were processed in cryoSPARC v.3 or v.4. A representative motion-corrected micrograph is shown at the top. Datasets were imported into cryoSPARC prior to motion correction and patch-based CTF estimation. Subsequent classification, ab inito and refinement steps are shown. The final model is colored by local resolution and presented at low contour (top) and high contour (bottom) values. Viewing angle distribution and GSFSC curves are shown after FSC auto-tightening for the final dimeric ternary complex density map.

**Supplemental Table 1:** Cryo-EM data collection, refinement and validation statistics.

|  | Monomeric ternary complex<br>(EMD-18809)<br>(PDB: 8R1F) | Dimeric ternary complex<br>(EMD-18810)<br>(PDB: 8R1G) |
| --- | --- | --- |
| **Data collection and processing** |  |  |
| Magnification | 120,000x | 120,000x |
| Voltage (kV) | 200 | 200 |
| Electron exposure (e<sup>-</sup>/Å<sup>2</sup>) | 50 | 50 |
| Defocus range (μm) | -0.2 to -2.0μm | -0.2 to -2.0μm |
| Pixel size (Å) | 0.84 | 0.84 |
| Symmetry imposed | C1 | C2 |
| Initial particle images (no.) | 16,800,000 | 11,300,000 |
| Final particle images (no.) | 297,000 | 85,396 |
| Map resolution (Å) | 3.67 | 3.99 |
| FSC threshold | 0.143 | 0.143 |
| Map resolution range (Å) | 3-11 | 3-11 |
| **Refinement** |  |  |
| Initial model used (PDB code) | 4XR8 | Monomeric ternary complex |
| Model resolution (Å) | 3.67 | 3.99 |
| FSC threshold | 0.143 | 0.143 |
| Map sharpening B factor (Å<sup>2</sup>) | 159.9 | 174.7 |
| **Model composition** |  |  |
| Non-hydrogen atoms | 8366 | 16732 |
| Protein residues | 1025 | 2050 |
| Ligands | 3 | 6 |
| B factors (Å<sup>2</sup>) | 213.38 | 120.08 |
| Protein | 213.35 | 120.07 |
| Ligand | 279.26 | 157.38 |
| **R.m.s. deviations** |  |  |
| Bond lengths (Å) | 0.008 | 0.006 |
| Bond angles (°) | 0.979 | 0.921 |
| **Validation** |  |  |
| MolProbity score | 0.69 | 0.91 |
| Clashscore | 0.36 | 1.35 |
| EMRinger score | 1.07 | 0.61 |
| Poor rotamers (%) | 0.21 | 0.27 |
| Ramachandran plot |  |  |
| Favored (%) | 97.74 | 97.84 |
| Allowed (%) | 2.26 | 2.16 |
| Disallowed (%) | 0.00 | 0.00 |

**Supplemental Table 2:** Molar mass measurements From SEC-MALS.

| Sample | Calculated MW (kDa) | Observed MW (kDa) | Possible oligomerization state |
| --- | --- | --- | --- |
| p53 | 43.65 (monomer) | 219 +/- 59 | Tetramer |
| E6AP-E6-p53 | 174.6 (1:1:1) | 671 +/- 175, 462 +/- 167, 221 +/- 74 | 4:4:4, 3:3:3, 2:2:0 or 2:0:0 |

## Methods

### Expression and purification of proteins

Full-length human E6AP, full-length p53, and human p53^DBD^, each with an N-terminal Strep tag, were cloned into pAC8 vectors for expression in insect cells. Using the Bac-to-Bac system (Life Technologies). recombinant baculoviruses were prepared in the Spodoptera frugiperda (sf9) cells. Full-length HPV16 E6, containing four amino acid subsitutions to improve solubility (C80S, C97S, C111S, C140S) was cloned into a pET-28(+) vector with an N-terminal 6xHis and SUMO tag for *Escherichia coli* expression.

E6AP was expressed in *Trichoplusia ni* High Five insect cells by infection of 25 ml of single baculoviruses per 1 L of High Five culture. Cells were harvested 36-48 hours after infection and lysed by sonication in buffer containing 25 mM Tris pH 8.0, 200 mM NaCl, 5% glycerol, 0.5 mM TCEP, 5 mM McCl2, 1x protease inhibitor cocktail (Roche Applied Science) and 0.1% Triton X-100. The lysate was clarified by ultracentrifugation at 40K RPM for 45 min. Supernatant was then loaded onto a gravity column containing a Strep-Tactin Sepharose bead slurry (IBA life sciences) for affinity chromatography. After binding, the column was washed with a low salt (200 mM NaCl), followed by high salt (1M NaCl), and then elute at 200 mM NaCl and 5 mM desthiobiotin. The Strep (II) eluted fractions were then diluted to 100 mM NaCl, prior to application on Q-column and then eluted with a linear gradient using 2M NaCl as Buffer B. The sample was then subjected to size exclusion chromatography (Superose 6, GE Healthcare) in a buffer contiaing 25 mM Tris pH 8.0, 150 mM NaCl, 5% glycerol, 0.5 mM TCEP. Pure fractions were collected, concentrated and sotred at −80°C. p53 and p53^DBD^ were likewise expressed in High Five insect cells and lysed in buffer containing 20 mM Tris pH 8.0, 1 M NaCl, 10% glycerol, 0.5 mM TCEP, 1x protease inhibitor cocktail (Roche Applied Science) and 0.1% Triton X-100 and underwent affinity chromatography as in E6AP. Strep (II) elutions were applied to a Heparin column, and eluted over a linear NaCl gradient, followed by size-exclusion (Superdex 200, Cytiva) with 150 mM NaCl. Pure fractions were collected, concentrated and stored at −80°C.

For HPV16 E6, 4-6 L of *E. coli* culture was inoculated in a 1:100 (v/v) ratio with an overnight pre-culture and incubated at 37°C with antibiotics. Upon reaching an OD_600 nm_ of 0.6-1, gene expression was induced with 0.5 mM IPTG induction. Cultures were allowed to grow overnight and harvested in the morning. Cells were harvested by centrifugation at 4°C for 10 min and resuspended in 100-200 mL lysis buffer containing 50 mM Tris pH 6.8, 400 mM NaCl, 20 mM Imidazole, 5% glycerol and 0.5 mM TCEP. The resuspended pellet was sonicated prior to lysate clarification through ultracentrifugation at 40K RPM for 1 hour. Filtered supernatant was loaded onto a HiTrap His column (Cytiva). After loading, the column was washed with wash buffer until absorbance is low and eluted with 10-80% containing 50 mM Tris pH 6.8, 400 mM NaCl, 500 mM Imidazole, 5% glycerol and 0.5 mM TCEP. The Eluted fractions were then further purified with a heparin column by first diluting to <200 mM NaCl before loading and using a linear NaCl salt gradient. Finally, samples were diluted/dialyzed to no less than 400 mM salt and flash frozen in 5-10% glycerol and stored at −80°C.

### Mass photometry

Before mass photometry measurements, protein dilutions were made in MP buffer (20 mM Tris-HCl pH 8.0, 100 mM KCl and 0.5 mM TCEP) and mixed in a 1:1:1 ratio (E6AP:E6:p53) and incubated at room temperature for 30 min. Data were acquired on a Refeyn OneMP mass photometer. First, 18 μl of MP buffer was added into the flow chamber followed by focus calibration. 2 μl of protein solution was then added to the chamber and movies of 60 or 90 seconds were aquired. Each sample was measured at least two times independently (n = 2) and acquired movies were processed and molecular masses were analysed using Refeyn Discover 2.3, based on a standard curve created with BSA and thyroglobulin.

### Analytical size-exclusion chromatography coupled to multi-angle light scattering (SEC-MALS)

Forty microliters of sample containing a mixture of p53, E6 and E6AP at concentrations of 11.25 uM each were injected onto a Superose 6 Increase 10/300 GL column (Cytiva) using an Agilent Infinity 1260 II HPLC system. In-line refractive index and light scattering measurements were performed using a Wyatt Optilab T-rEX refractive index detector and a Wyatt miniDAWN TREOS 3 light scattering detector. System control and analysis was carried out using the Wyatt Astra 7.3.1 software. System performance was checked with BSA (initial and final run).

### Cryo-EM

E6AP, E6 and p53 proteins at ∼ 2mg/ml in a 1:1:1 stoichiometry were applied to a 10-30 % sucrose gradient composed of 20 mM HEPES ph 8.0, 100 mM KCl, 0.5 mM MgCl2, and 0.5 mM TCEP and 0-0.2% (w/v) glutaraldehyde or 0-2 mM bissulfosuccinimidyl suberate according to the “GraFix” method^20^. Gradients underwent ultracentrifugation at 37K RPM for 17 hours in a SW-60 swinging bucket rotor. Gradients were harvested by piston fractionation (BioComp) with peak fractions found by absorbance traces or SDS-PAGE. Peak fractions were pooled and underwent overnight dialysis to remove sucrose, prior to concentrating to ∼30μL using Amicon centrifugal filter concentrators (Merck-Millipore).

2.5μL of ∼1.2 mg/ml sample were applied to UltraAuFoil 1.2/1.3 300 mesh (Quantifoil) or 1.2/1.3 Cu 300-mesh grids (Quantifoil) with or without graphene oxide deposited on the grid surface that were freshly glow dishcharged with a Pelco EasyGlow (15 mA current, 45 seconds). For the GA crosslinked datasets, additional grids were prepared with the following added detergents with or without graphene oxide deposited on the grid surface: 0.068% octyl-β-glucoside, 0.005% NP40, 0.05% LMNG, 0.05% CHAPS, 0.005% DDM and 2mM fluorinated fos-choline. Grids were plunge-freezed into liquid ethane following a 3-4 second blot time at 95% humidity and 4°C using a Vitrobot Mark IV (FEI).

Automatic data collection was done using EPU 3.0 (Thermo Fisher Scientific) on a Glacios (Thermo Fisher Scientific) electron microscope operating at 200 kV. Images were recorded using a Falcon 4 direct electron detector (Thermo Fisher Scientific) using a nominal magnification of 120,000x. All datasets were recorded with an accumulated total dose of 50 e^−^/Å^2^ fractionated into 50 frames. The targeted defocus values ranged from −0.2 to −2.0μm.

Real-time evaluation was performed using CryoFlare1.10^38^ during acquisition. Images were motion corrected using Relion3 motioncorr implementation^39^ by applying a dose-weighting scheme to generated a motion-corrected sum of all frames. CTF estimation was done using the patch CTF implementation in cryoSPARC v.3 or v4^40^. Particles were picked using either blob picker or template picker in cryoSPARC v.3 or v.4^40^. Each dataset was further processed in cryoSPARC v.3 or v.4^40^ which included 2D and 3D classification, 3D refinement and CTF refinement. Resolution values reported for each reconstruction were calculated based on the gold-standard Fourier shell correlation curve (FSC) at 0.143 criterion^41^, correcting for the effects of soft masks using high-resolution noise substitution^42^. Local resolutions for each final map was estimated using MonoRes (XMIPP) implementation in cryoSPARC v.3 or v.4^43^.

### Model building and refinement

The structure of full-length E6AP was first predicted with Alphafold2^21^ and docked into the map of the monomeric ternary complex, along with E6 and p53 chains from the previous crystal structure of the ternary complex (PDB: 4XR8)^19^ using the ChimeraX fit-in-map^44^. Restrained flexible-fitting was done with ISOLDE^45^, and local corrections were done with Coot^46^ and ISOLDE. B-factor fitting and coordinate-constrained minimization was done with Phenix real-space refine^47^. The dimeric ternary complex model was built by docking two copies of the monomeric complex model into the dimeric map using ChimeraX fit-in-map, followed by local corrections done in ISOLDE and B-factor fitting using Phenix real-space refine. Validation was carried out with Phenix, Molprobity^48^, and EMRinger^49^. Figures were prepared with ChimeraX.

